# Erasable Serum Markers

**DOI:** 10.1101/2025.05.08.652140

**Authors:** Shirin Nouraein, Sangsin Lee, Honghao Li, Vidal A. Saenz, Emma K. Raisely, Vincent P. Costa, Jerzy O. Szablowski

## Abstract

Gene expression in the brain is typically evaluated using invasive biopsy or postmortem histology. Serum markers provide an alternative way to monitor the brain, but relatively few such markers exist. Additionally, the origin of serum markers often cannot be localized to a specific cell population, and monitoring dynamic changes in their gene expression is compromised by the same factor that makes the markers detectable – long serum half-life. Here we propose a paradigm to improve the sensitivity of serum marker measurement by modifying the markers *in vivo*, called erasable serum markers, or ESM. As a proof of concept, we use a well-controlled system with known half-life and tunable serum levels. This system, released markers of activity, or RMAs enable measurement of transgene expression in the brain through a simple blood test. RMAs are stable in blood, with a half-life of >100 h and can detect expression from as few as 12 neurons in mice. However, their long serum half-life also generates long-lasting background signals when RMA are used to track temporal changes in gene expression. By engineering on-demand erasable RMAs and injecting an intravenous targeted protease, we reduced RMA background signal by more than an order of magnitude without compromising the detection sensitivity. Similarly to previous RMA iteration, our approach showed a 65,000-fold increase in their signal over the baseline when expressed in a single brain region. Furthermore, we demonstrated that this erasable RMA system improves the dynamic range of detection for low-level promoter activity that is driven by physiological levels of c-Fos.

## INTRODUCTION

Serum markers are vital tools in diagnostic medicine and *in vivo* research and can provide insights into the biological and physiological functions of living organisms. The determination of serum marker levels entails a simple blood collection procedure^1^, which enables longitudinal studies, where the same individual can be studied over time^2^. However, naturally occurring serum markers can only provide information on a selected subset aspects of bodily functions. Additionally, serum markers often lack the sensitivity or characteristics that would be optimal for their detection. For example, long half-life of the serum markers in the blood is beneficial to enhance their detectability, but also leaves background levels of markers that hamper tracking dynamic changes in these markers’ production rate. While the natural marker properties are usually accepted as they are with all their pros and cons, we propose an alternative paradigm. By modifying the markers in vivo, it may be possible to tune their properties to improve measurement. For example, with a proteolytic cleavage of specific markers in blood, it may be possible to erase the background levels of these markers before a measurement is needed, forming erasable serum markers (**ESM**). Such cleavage could then be followed by measurement of markers at multiple timepoints to determine the rate of their replenishment, and thus the rate of markers’ production. As a proof of concept of this paradigm, we decided to use a well-controlled system of synthetic serum markers called released markers of activity (**RMA**). RMAs are genetically-encoded reporters that can be expressed in the brain through gene therapy and secreted across the blood–brain barrier (BBB)^3,4, 5^ into the circulation^6^. RMA can be used to monitor the expression of specific genes in the brain by placing them under the control of a specific promoter^6^ or RNA-based sensor^7^. Specifically, RMA were shown to be useful in observing neuronal activation by monitoring c-Fos^6,7^ or Arc expression^7^. Additionally, RMA can be detected with high sensitivity from as few as 12 neurons in the mouse brain with 5 µL of blood^6^, and their production rate can be tuned by tying them to various promoters.

The detection of RMA at a high degree of sensitivity can be attributed to three features. First, RMA are released into the blood, which makes them easily accessible. Second, they can be paired with an easily detectable marker, such as luciferase. Third, they have a long half-life in serum (∼100 hours), allowing them to accumulate in blood over time. However, this long half-life also results in the retention of the RMA signal, leading to a sustained background signal.

Dynamic changes in gene expression can be discerned by measuring the rate of change in serum RMA levels over multiple timepoints, rather than overall RMA levels, decoupling the half-life of RMAs from their temporal resolution. High background signal, however, limits the dynamic range of measurement. Although the background signal could be lowered by simply reducing the half-life of RMA, such a reduction would lower the overall signal level and the sensitivity of measurement. Additionally, the Fc region of the immunoglobulin G (IgG) segment in RMA contributes to their long half-life and facilitates their transfer across the BBB. Therefore, modifications to the Fc region can alter the reverse transcytosis of RMA across the BBB and their half-life in serum.

Herein, we present an innovative design solution that enables efficient reverse transcytosis of RMA across the BBB and achieves a more than 10-fold reduction in the RMA background signal without compromising on detection sensitivity. To overcome the drawback of the prolonged half-life of RMA, we introduced a protease cleavage site within the RMA molecule between the luciferase tag and the Fc region of IgG. We call this modified markers fast-erasable RMA (**feRMA**). Upon *in vivo* cleavage with tobacco etch virus (TEV) protease^8-13^, the Fc region separates from luciferase, leading to the rapid clearance of the remainder of feRMA from the serum. Consequently, feRMAs can cross the BBB owing to the Fc region but are rapidly cleared from the serum on-demand following the simple injection of a protease. Our results demonstrated that over 90% of the feRMA protein was degraded following the expression of TEV protease both *in vitro* and when secreted from transduced cells. We further explored the kinetics of feRMA expression after delivery using an adeno-associated virus (AAV) as well as the potential of feRMA to monitor gene expression in the brain over multiple timepoints. We used c-Fos sensing through a robust activity marking (RAM) system^14^ to demonstrate the ability of feRMA to record baseline levels of c-Fos to show sensitivity of this measurement. Overall, the results showed that feRMA enables erasable, repeatable reduction of the RMA background signal and longitudinal monitoring of targeted gene expression in the brain.

## RESULTS

### Developing an erasable synthetic serum marker

The feRMA reporter is comprised of three main parts: 1) a secretion tag for exporting the protein outside the cells; 2) an easily detectable tag, such as luciferase; and 3) an IgG Fc region that allows reverse transcytosis of the reporter from the brain to the blood. The feRMA design differs from the RMA design in the inclusion of a protease cleavage site between the luciferase marker and the Fc domain (**Fig. 1a**). Given that the extended half-life of RMA is attributed to the Fc region^6,15^, we posited that the cleavage of the Fc region should reduce the half-life of free luciferase, thereby reducing the background signal. However, because feRMA within the brain are protected from cleavage by the BBB, they can still transcytose into the blood without being modified by the protease. As a result, this feRMA design ensures efficient transcytosis, a long half-life when sensitivity is required, and a shorter half-life on demand achieved by the intravenous injection of TEV protease.

**Figure 1:**
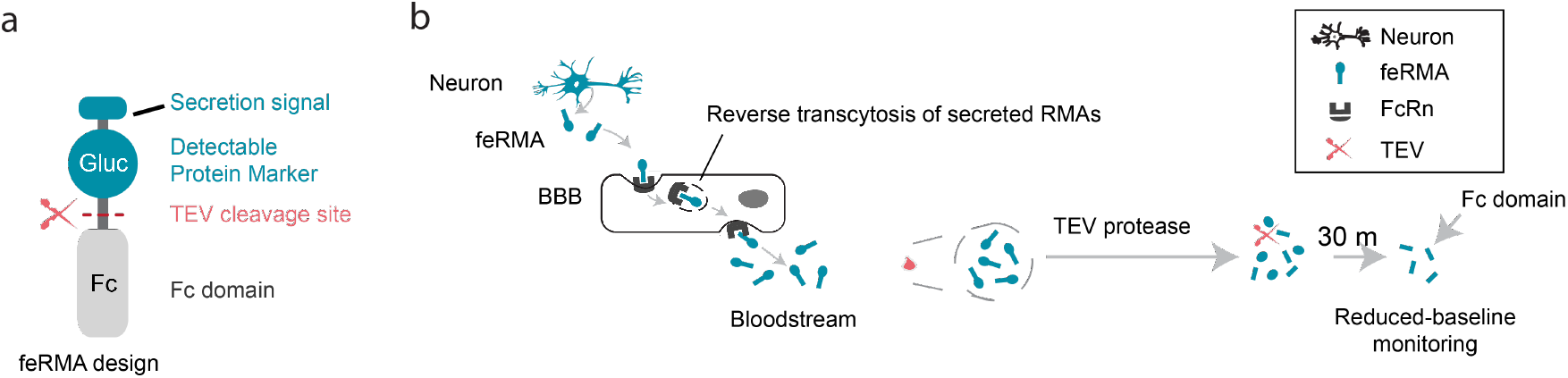
Schematic depicting the mechanism of fast-erasable released markers of activity (feRMA). **a**, feRMA comprises three main components: (i) a secretion tag for exporting the protein outside the cells; (ii) a detectable protein marker such as Gluc; and (iii) Fc, a reverse transcytosis domain that also extends the half-life of the molecule in blood. Components (ii) and (iii) are linked by a TEV cleavage site, making the protein specifically cleavable by TEV protease. Cleavage of feRMA using TEV protease leads to rapid serum clearance and signal reduction due to the detachment of Gluc from the long-lasting Fc domain. **b**, feRMA are genetically encoded proteins capable of crossing the blood–brain barrier (BBB) and entering the bloodstream, where they can be erased using TEV protease or detected through a blood test.

We used a sequence-specific TEV protease in this study^8,9^. TEV protease is commonly used in protein purification protocols to separate proteins from affinity tags^10-12^. Although TEV protease activity has not previously been validated in circulation, it has been successfully expressed as a transgene *in vivo*^12,13^. Therefore, TEV protease presented as a feasible proof-of-concept protease for engineering erasable serum markers.

We developed several feRMA constructs by introducing linkers of different lengths between the TEV protease cleavage site and the remainder of the feRMA protein (**Supplementary Table 1**). To confirm that TEV protease can cleave each feRMA construct, 480 pmols of each feRMA construct was incubated for 1 h with 370 pmols of TEV protease. Subsequent analysis using sodium dodecyl sulfate–polyacrylamide gel electrophoresis revealed the presence of an feRMA band (44 kDa) after the incubation of each construct with TEV protease (Fig. 2a). The cleavage efficiencies of feRMA1, feRMA2, and feRMA3 were 92.62% ± 1.06%, 95.04% ± 1.27%, and 94.6% ± 0.92%, respectively (n = 4 each group, p = 0.4008, one-way analysis of variance [ANOVA], mean and standard error of the mean [SEM] provided; Fig. 2a, b). We also assessed the ability of feRMA to be released from transfected cells. We expressed three Gluc-TEVcs-Fc constructs under the neuron-specific hSyn promoter in PC-12 cells, a widely used murine cell line for studying neurosecretion^16^. Culture media were collected at 120 h post transfection, and the relative amount of secreted Gluc-TEVcs-Fc in the media was measured using a luciferase assay (Fig. 1c). All three feRMA constructs were detected in the media. feRMA2 and feRMA3 exhibited a higher luciferase signal than feRMA1. No significant difference was observed between the expression levels of feRMA2 and the regular RMA protein marker (Fig. 1c). These results indicate that the designed feRMA constructs can be effectively cleaved by TEV protease and expressed in mammalian cells.

**Figure 2:**
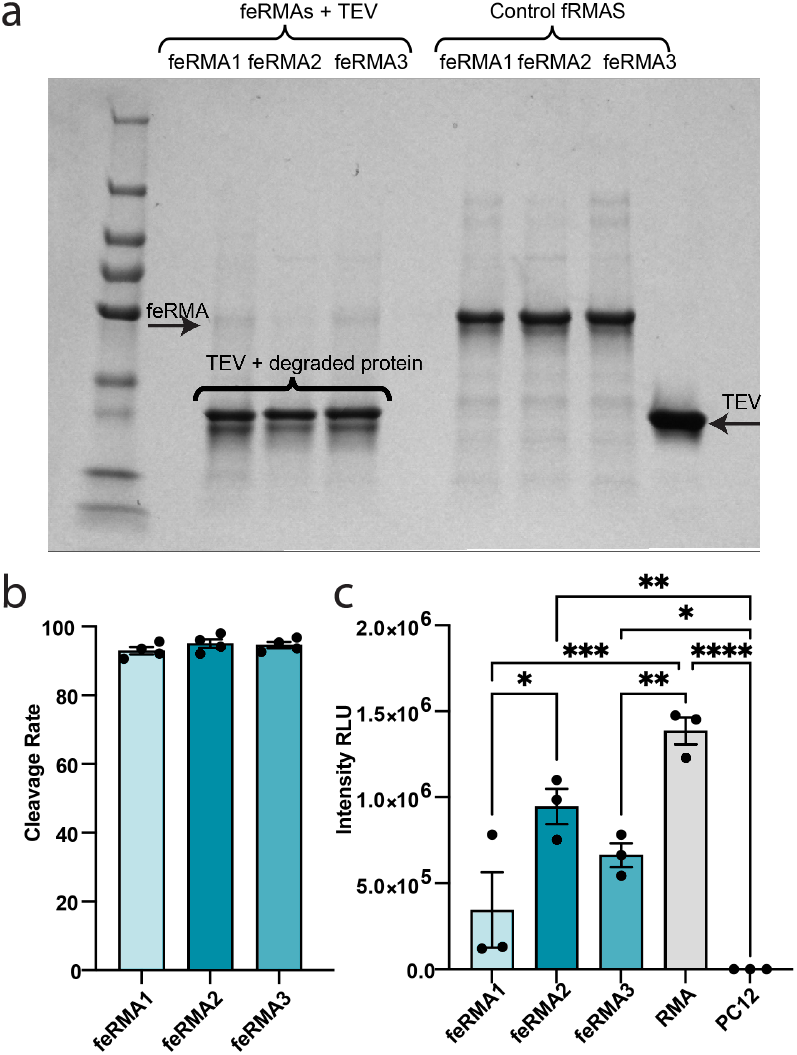
*In vitro* cleavage of feRMA, **a**, SDS–PAGE testing the cleavage of feRMA proteins with TEV protease for 1 h at 37°C. Product concentrations were calculated from the intensities of the bands. **b**, Results indicated an average cleavage efficiency of 92.96% ± 1.06%, 95.04% ± 1.27%, and 94.61% ± 0.92% for feRMA1, feRMA2, and feRMA3, respectively (n = 4 each group, F _2,9_ = 1.014, p = 0.4008, one-way ANOVA), upon exposure to TEV protease. **c**, feRMA secretion by PC-12 cells at 120 h post transfection. Relative luminosity unit (RLU) values measured from the culture media revealed the signal peptide–dependent secretion of Gluc-TEV_CS_-Fc RMA constructs. n = 3 independent cultures were analyzed. In comparison with the signal at different time points, using two-way ANOVA, with Tukey’s test (F_4.8_ = 26.35, p = 0.0001, P = 0.0951 (feRMA2 vs. RMA), P = 0.0007 (feRMA1 vs. RMA), P = 0.0076 (feRMA3 vs. RMA). Means and standard errors of the mean (SEM) are presented in bar graphs. Non significant comparisons not shown for clarity.

### feRMA can be secreted from the brain into the blood

To assess the translocation of feRMA proteins from the brain into the blood, we directly injected feRMA proteins (20 pmols) into the caudate/putamen (CP) of mouse brains. We expected the feRMA proteins to reverse-transcytose across the BBB, a process mediated by the interaction between the Fc domain and the neonatal Fc receptor^17^, as demonstrated in our previous studies^6,7^ (**Fig. 3a**). Blood was collected at 2, 6, 12 (for feRMA2 and feRMA3), and 24 h post injection. Within 6 h post injection, the plasma concentrations of all three Gluc-feRMA constructs significantly increased (feRMA1 ****p < 0.0001, feRMA2 **p = 0.0013, and feRMA3 **p = 0.0059 one-way ANOVA; **Supplementary Fig.1**). The mice were euthanized after 24 h through cardiac perfusion. Their brains were sectioned for histology and immunostained with anti-Gluc antibody. Gluc-feRMA was not detected in the brain sections, suggesting the complete export of feRMA proteins from the brain (**Supplementary Fig.1**). Further, although all three feRMA protein constructs successfully crossed the BBB, feRMA2 exhibited a significantly higher luciferase signal in the bloodstream at 6 h compared with the other constructs (***p = 0.001 for feRMA1 vs. feRMA2 and ***p = 0.0004 for feRMA2 vs. feRMA3, n = 4, F_2,36_ = 29.84, p < 0.0001, two-way ANOVA, means and SEM provided; **Fig. 3b**). Based on these results and those of our *in vitro* tests (**Fig. 2**), we chose feRMA2 for further experiments.

**Figure 3.**
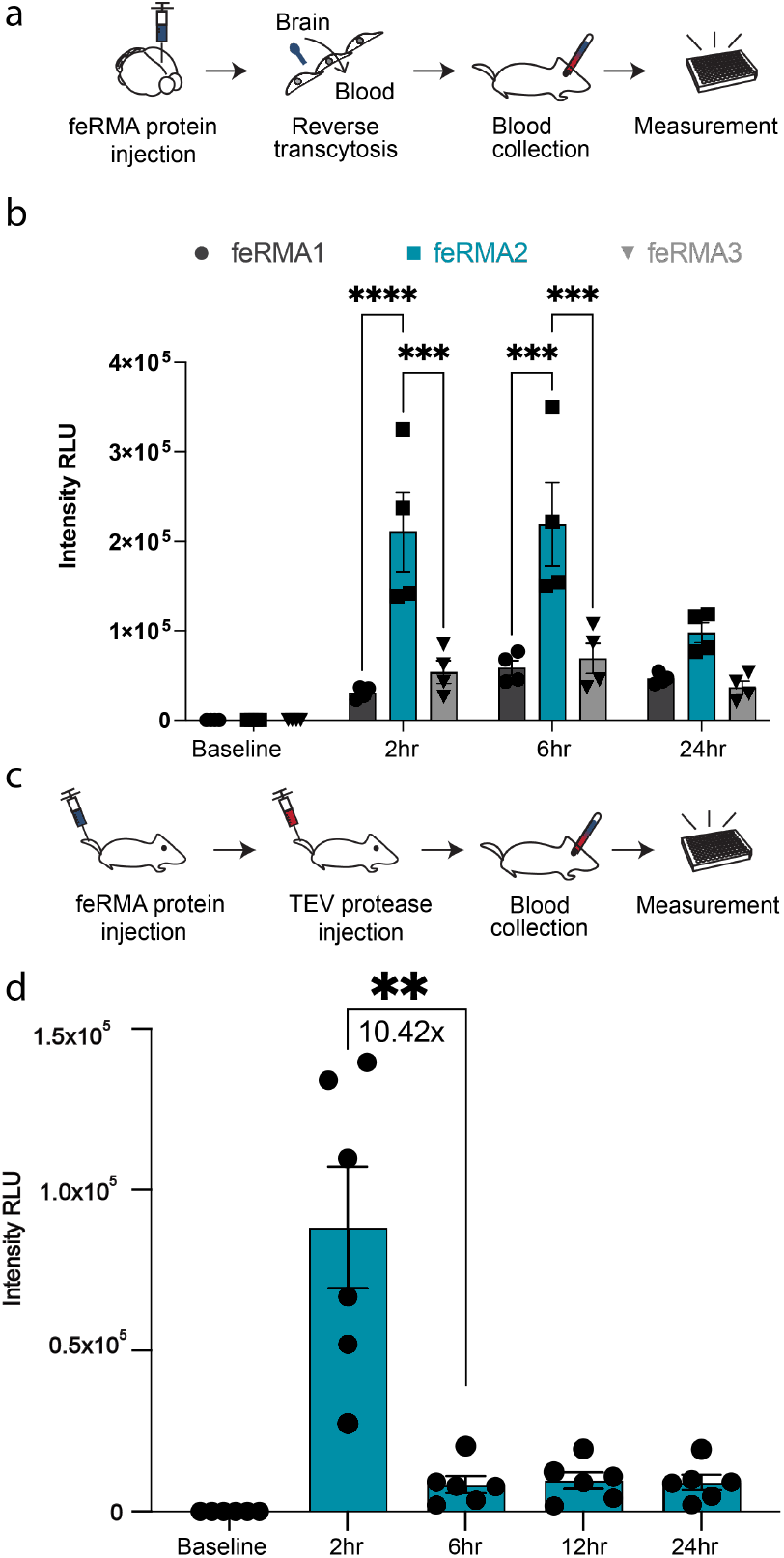
Brain-to-blood transcytosis of the TEV protease– cleavable feRMA reporter. **a**, Experimental scheme for testing the reverse transcytosis of feRMA across the BBB into the bloodstream. **b**, Plasma concentration of feRMA measured from the collected blood after injecting 20 pmols of feRMA constructs into the caudate/putamen of mice brains. N = 4 mice per group were analyzed. Two-way ANOVA for comparing all three constructs at different timepoints (***p = 0.001 for feRMA1 vs. feRMA2 and ***p = 0.0004 for feRMA2 vs. feRMA3 at 6 h. P-values obtained using two-way ANOVA [F2,36 = 29.84, p < 0.0001] with Tukey’s test. **p < 0.01, ***p < 0.001. ****p < 0.0001). c, Experimental scheme for testing the in vivo cleavage of feRMA. d, Plasma concentration of feRMA at 2 h following the injection of 12 nmols of TEV protease. N = 6 independent mice were analyzed. p = 0.0052 according to one-way ANOVA (F1.017,5.086 = 21.84) using Sidak’s test for comparing the fold-change in the signal at 2 h with that at 6 h. Means with SEM values presented in the bar graphs.

Next, we evaluated the cleavage of the feRMA protein *in vivo* following an intravenous injection of TEV protease (**Fig. 3c**). Because the activity of TEV protease has not been tested *in vivo*, we started with a low dose. We administered a vehicle and two doses of TEV protease (12 pmols and 12 nmols) through the tail vein. The feRMA dose was maintained at 20 pmols. The vehicle control group showed a significant drop in feRMA levels (1.34 ± 0.17 fold, p = 0.0022, one-way ANOVA, means and SEM provided; **Supplementary Fig. 2b**) owing to the clearance of the protein from the blood^6^. Interestingly, treatment with 12 pmols of TEV protease did not significantly decrease feRMA levels (p = 0.8844, one-way ANOVA, means and SEM provided; **Supplementary Fig. 2b**); however, treatment with 12 nmols of TEV protease significantly reduced feRMA levels 4 h after protease injection (10.42 ± 0.21 fold, p = 0.0052, means and SEM provided, one-way ANOVA; **Fig. 3d**).

### Tracking gene expression in the brain using feRMA

We hypothesized that feRMA can be used to track transgene expression in the brain using a blood test. To test this hypothesis, we encoded the feRMA construct in PHP.eB, a BBB-permeable AAV^18^, under a neuron-specific human synapsin (hSyn) promoter^19^. In addition, we placed the coding sequence of green fluorescent protein (GFP) downstream of an internal ribosome entry site (IRES) to allow the facile quantification of transduction efficiency, a strategy employed in our previous study^6^. We injected 2.3 × 10^9^ (200 nL) of viral particles (VP) per g of body weight encoding feRMA directly into the brains of mice divided into three groups according to the site of injection: CP of the striatum, cornu ammonis of the hippocampus (CA1), or substantia nigra (SN) in the midbrain region.

We collected blood samples from each mouse at five time points, namely, baseline, 1 week after AAV injection, 30 min after TEV protease injection at the end of week 1 (Week 1 + TEV), 3 weeks after AAV injection, and 30 min after TEV protease injection at the end of week 3 (Week 3 + TEV; **Fig. 4a**). All blood samples were frozen after collection and simultaneously subjected to the luciferase assay.

**Figure 4.**
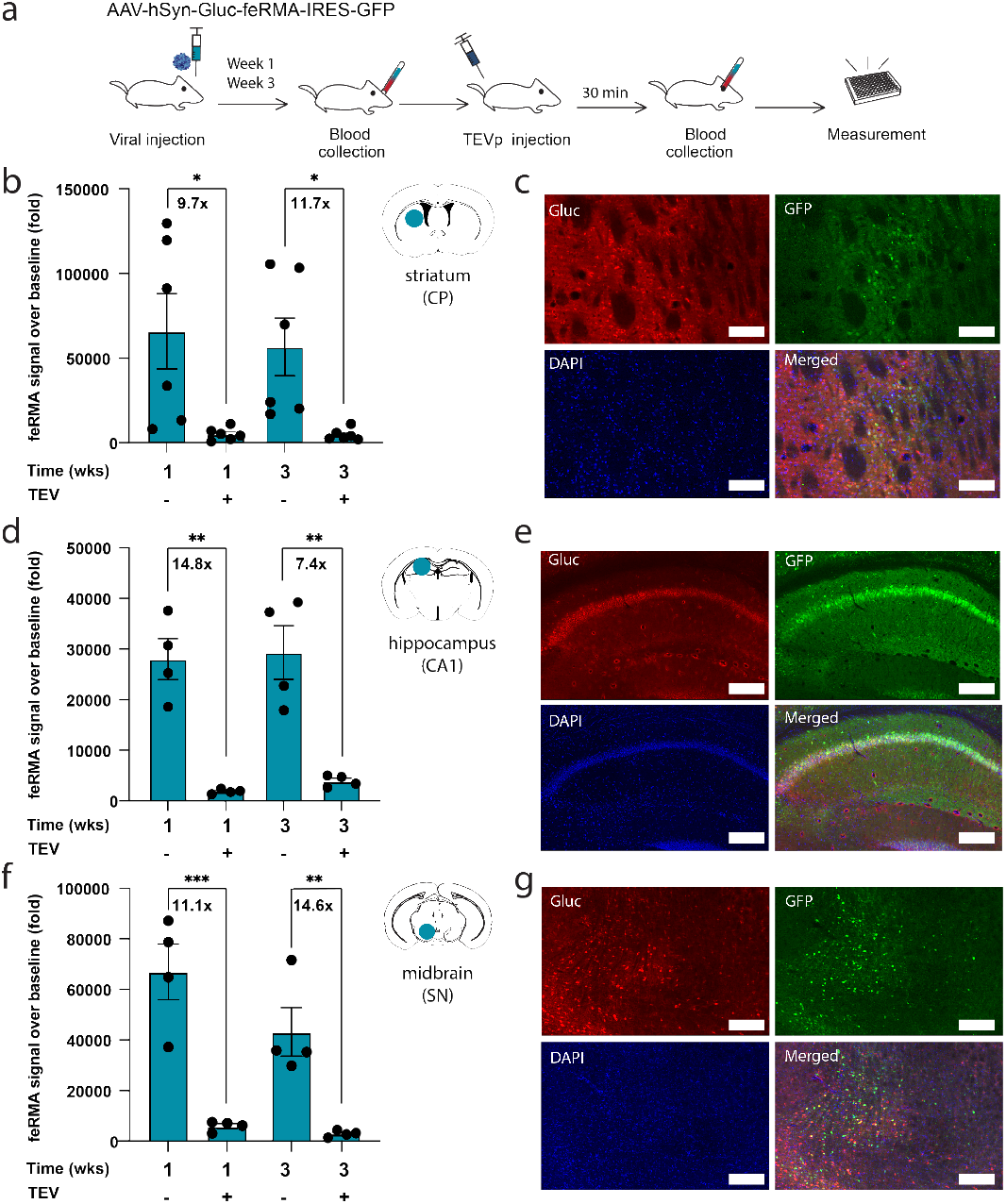
feRMA provides an erasable signal that enables the monitoring of gene expression in multiple brain regions. **a**, Experimental scheme for detecting gene expression in the brain using feRMA. Blood was collected from mice to establish the baseline luminescence level. Then, AAVs encoding Gluc-feRMA were injected into the striatum, hippocampus, or midbrain of the mice. At 1–3 weeks after injection, blood was collected to measure the Gluc-feRMA signal. Immediately afterward, TEV protease was injected intravenously, and blood was collected 30 min later to verify the erasure of feRMA. **b**, feRMA levels were normalized to the pre-AAV injection baseline, shown before and after TEV protease administration (striatum group, n = 6). **c**, Representative images of the striatal expression of feRMA (red), expression of GFP mediated by an internal ribosome entry site downstream of feRMA (green), and a nuclear co-stain (DAPI, blue) at the site of delivery. **d, e**, Replication of the experiment described in panels b and c with feRMA expression in the CA1 region of the hippocampus (n = 4) and **f, g**, the substantia nigra of the midbrain (n = 4). Blue circles indicate the target injection sites. The numbers between each bar refer to the fold change in feRMA signal following injection of TEV protease. Overall effects of TEV on the feRMA signal was assessed using a mixed-effects ANOVA for CP (*F*(1,3) = 9.6001, *p* = 0.0269), CA1 (*F*(1,3) = 46.8714, *p* = 0.0064), and SN *F*(1,3) = 39.7524, *p* = 0.0086). Pairwise comparisons assessed effects of TEV protease administration at 1 and 3 weeks, respectively: CP (*p* = 0.019 and *p* = 0.0347; panel b), CA1 (*p* = 0.0058 and *p* = 0.0065; panel d), and SN (*p* = 0.008 and *p* = 0.0061; panel f). Means with SEM presented in bar graphs. Right: Gluc-feRMA expression at the local injected sites. Scale bars, 100 µm.

The luciferase assay confirmed protein expression as early as at week 1 following gene delivery in all three experimental groups. Compared with the baseline, the luciferase signal increased by 65,524 ± 20,027 fold, 28,308 ± 4,066 fold, and 63,510 ± 10,382 fold in the CP, CA1, and substantia nigra groups, respectively (means ± SEM). The feRMA levels significantly decreased in all the three groups (7.4 - 14.8-fold) at 30 min post TEV protease injection, as evidenced by a reduction in the background signal (**Fig. 4b–g**).

After 3 weeks, the brains were harvested and sectioned for the histological analysis of feRMA transduction. GFP was used as a more robust tissue marker to monitor transduction instead of Gluc because of the possibility of Gluc-feRMA transcytosing out of the brain into the blood, thereby reducing detectable Gluc protein in the brain. All three targeted regions displayed significantly more GFP-transduced cells compared with contralateral controls, confirming the success of AAV delivery (**Supplementary Fig. 3b–d**).

Overall, our results demonstrate that intravenous administration of TEV protease cleared most of the feRMA molecules from the blood within 30 min of injection at two different timepoints. Moreover, the designed feRMA constructs could successfully monitor transduction in the brain, with signal levels comparable to those obtained using RMA^6^.

### feRMA improves the dynamic range for measuring baseline c-Fos expression

Minimally invasive monitoring with RMA is particularly useful for observing long-term changes in gene expression. We hypothesized that reducing the background signal using feRMA will enable more effective measurement of gene expression in mice over multiple timepoints. Specifically, given that RMA used in previous studies could only monitor strong neuronal activation with chemogenetic activation^6,15^, we were interested in determining whether feRMA could accumulate over time when monitoring low baseline levels of c-Fos activity. Accordingly, we designed a proof-of-concept experiment evaluating feRMA expression in response to baseline neuronal activity in the brain. To gate the recording to a specific time frame, we used the RAM system^14^—a doxycycline (Dox)-dependent Tet-Off system—to couple the feRMA reporter gene to a synthetic Fos promoter. In the RAM system, the presence of Dox suppresses the expression of downstream proteins, whereas its withdrawal activates protein expression in response to c-Fos promoter activity^14,20.^ We prepared two AAV-PHP.eB constructs: one encoding the hM3Dq and RAM system under the hSyn promoter and the other encoding the feRMA system under the tetracycline response element (TRE) promoter (**Fig. 5b**). Both constructs were co-injected into the CA1 region of the hippocampus in mice. The mice were initially fed a Dox-containing diet starting at 48 h before virus injection. This diet was replaced with a Dox-free diet at 72 h before the beginning of recording. Blood samples were collected at five timepoints: immediately after Dox withdrawal (baseline) and at 48, 48.5, 96.5, and 97 h. Half of the mice received vehicle (**Fig. 5c**), while the other half received TEV protease at 48 and 96.5 h following Dox withdrawal to evaluate the erasure of the feRMA signal (**Fig. 5d**).

**Figure 5:**
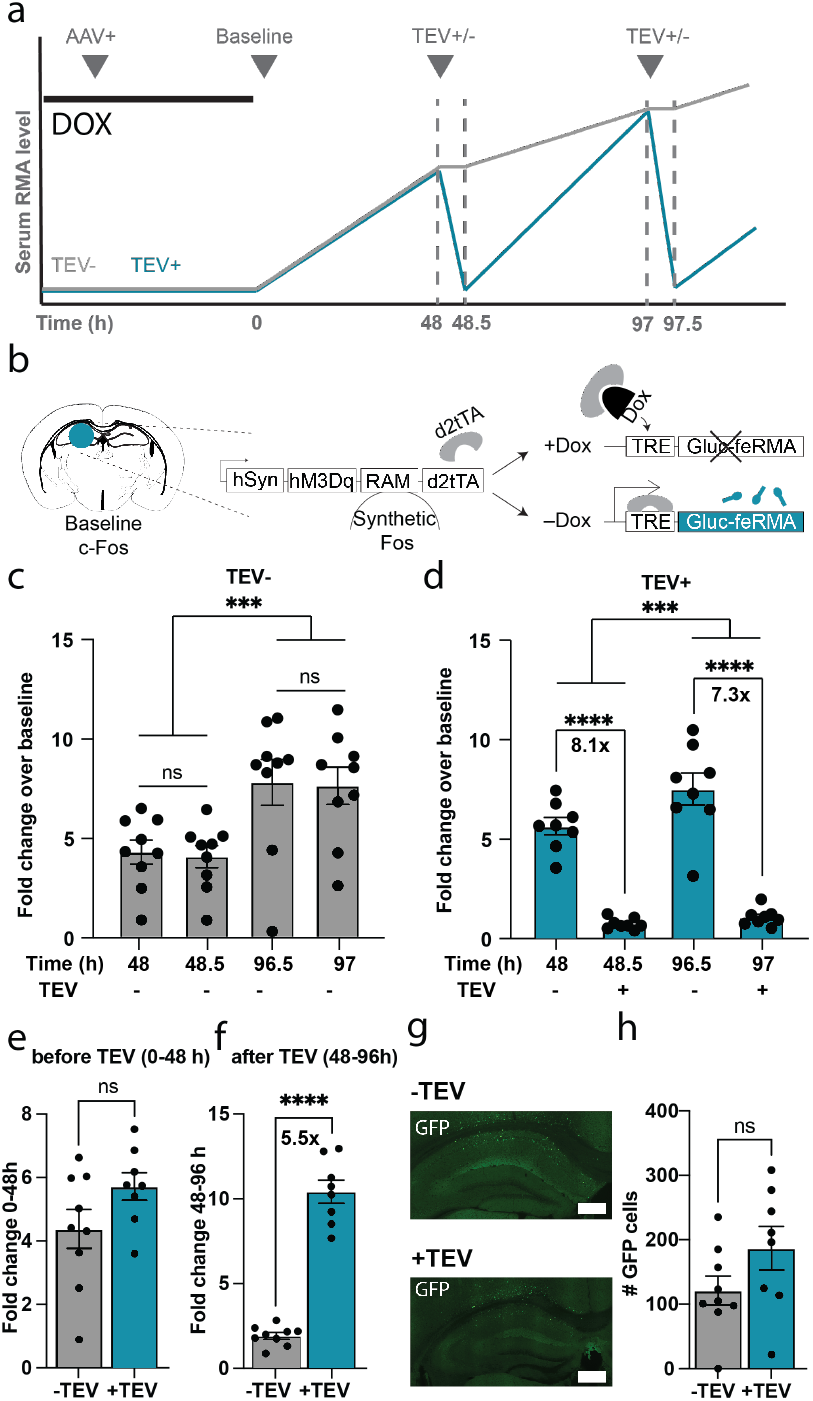
feRMA approach improves the dynamic range of measurement of baseline c-Fos expression. **a**, Experimental timeline for the detection of baseline c-Fos expression in the hippocampus using feRMA. Initially, the mice received a Dox-containing diet to suppress the production of RMA until the time of observation. After **48 h**, the mice received an intraparenchymal brain injection of AAVs to express feRMA reporter genes in the hippocampus. A week later, Dox was with-drawn and feRMA was allowed to accumulate in the serum for 48 h to measure the average baseline c-Fos activity in the hippocampus. A blood sample was collected, TEV protease or vehicle was administered intravenously, and a second blood sample was collected 30 min later. The experiment was repeated 48 h later using the same animals to demonstrate the repeatability of the measurement. **b**, To record c-Fos activity, we used the robust activity marking (RAM) system. In the RAM system, a tetracycline transactivator (d2tTA) is expressed, which drives the expression of Gluc-feRMA in the absence of Dox. AAV was injected into the CA1 region of the hippocampus. **c**, In the absence of TEV protease, the feRMA signals did not differ significantly between the 48- and 96-h samples or between the repeated blood samples collected 30 min later (F(1,8) = 0.2387, p = 0.6396). However, feRMA levels significantly increased in the serum between 0 and 48 h and between 48 and 96.5 h after Dox withdrawal (F(1,8) = 34.988, p = 0.0004). **d**, Administration of TEV protease caused a significant reduction in RMA levels within 30 min (F(1,7) = 14.5108, p = 0.0066), with significant differences observed at both the 48- and 96.5-h timepoints (48: *F*(1,9) = 83.8631, *p* < 0.0001; 96.5: *F*(1,9) = 144.169, *p* < 0.0001). feRMA also accumulated over time (*F*(1,7) = 22.4108, *p* = 0.0004). **e**, Before the administration of TEV protease, the average rate of accumulation of the feRMA signal over 48 h was comparable between the vehicle and TEV protease groups (p = 0.098, two-tailed unpaired t-test). **f**, Administration of TEV protease suppressed the baseline signal and increased the dynamic range of measurement over 48 h of RMA accumulation. The TEV protease– treated group showed a significantly higher fold-change increase in RMA signal over the next 48 h (p < 0.0001, two-tailed unpaired t-test). **g**, Representative images of GFP expression in both vehicle and TEV protease–treated groups **h**, GFP expression in the RAM system is driven by the same c-Fos-dependent circuit as feRMA. Thus, the cellular GFP accumulation is indicative of average c-Fos activity in the cells and can be regarded as a longer-lasting proxy of c-Fos expression over time. We found that GFP levels were comparable between the vehicle and TEV protease–treated groups of mice. ns, p > 0.05; *, p < 0.05; **, p < 0.01; ***, p < 0.001. ****m p < 0.0001. Panels c and d evaluated effects using repeated-measures ANOVA. Panels e and f used two-tailed between group t-tests. Means with SEM presented in bar graphs.

In the absence of TEV protease, there was an equivalent increase in the feRMA signals measured 30 minutes apart after 48 and 96 h (*F*(1,8) = 0.2367, *p* = 0.6396) and the feRMA signal at 96 h was significantly higher than that at 48 h (*F*(1.8) = 34.9880, *p* = 0.0004). This contrasted with the effects of injecting the TEV protease. At both 48 h (F(1,9) = 93.8631, *p* < .0001) and 96 h (*F*(1,9) = 144.169, *p* < .0001), the TEV protease injection reduced the feRMA signals relative to signals measured 30 minutes prior to TEV protease injection (*F*(1,7) = 130.1980, *p* < 0.0001). Despite, the TEV protease mediated reduction in the feRMA signal at 48 h the feRMA signal continued to increase and was larger at 96 h than at 48 h (F(1,7) = 22.4108, *p* = 0.0004). Although, the fold increase in the feRMA signal between 48 and 96 h was smaller in the group administered TEV (1.52 fold) vs. vehicle (1.82 fold).

Next, we assessed whether the injection of TEV protease increases the rate of signal accumulation over time. We found that before the injection of TEV protease (0–48 h after Dox withdrawal), the vehicle and TEV protease groups did not differ significantly in terms of the feRMA signal (p = 0.057, 1.3-fold, **Fig. 5e**). However, in the next 48 h following the injection of TEV protease or vehicle, the feRMA signal in the TEV protease group significantly increased by 5.5-fold (p < 0.0001, two-tailed unpaired t-test, **Fig. 5f**).

To verify that the transduction and neuronal activity were comparable between the experimental groups, the brain sections were immunostained with GFP. As expected, we observed no significant difference in the expression of intracellular GFP between the TEV+ and TEV− groups (p = 0.115, two-tailed unpaired t-test, **Fig. 5g–h**).

## DISCUSSION

Recent advancements in protein engineering have enabled the monitoring of gene expression in the brain^21^. However, most of the newly developed approaches face challenges in sensitivity and the specific targeting of brain regions^22^. Bioluminescent imaging is a promising candidate for monitoring gene expression; however, its effectiveness is limited by attenuation of signal through the skull and the need for bioluminescent substrates to cross the BBB^23^. RMA have emerged as a sensitive tool for the noninvasive monitoring of gene expression in the brain. The sensitivity of RMA is partly attributed to their long serum half-life^6^. However, their long half-life limits the dynamic range of measurement during the temporal monitoring of changes in gene expression. The presence of long-lasting background levels of RMA leads to blunted responses to changes in gene expression. To address this limitation without compromising the sensitivity of RMA, we engineered a new marker called feRMA by modifying the original RMA to reduce its half-life on demand. This was achieved by inserting a TEV protease cleavage site to separate the Gluc detection label from the RMA domain, enabling the reverse transcytosis of the marker and its stabilization within the serum^6,24,25^.

The results of our tissue culture experiments demonstrated that all the designed Gluc-TEVcs-Fc variants accumulated in the culture media over time, with no significant differences between the levels of the variants and those of the main RMA marker^26^. This indicated that fusing TEV_CS_ to Gluc-Fc does not compromise the ability of the marker to be secreted. Furthermore, we observed that all the variants could cross the BBB and enter the bloodstream, as evidenced by the bioluminescence signals detected in mice serum. All the variants successfully crossed the BBB, and this reverse transcytosis occurred within 2 h of intracranial injection.

Upon injection of the marker in the CP of the striatum, feRMA signals were significantly elevated compared with the baseline. Notably, the feRMA2 construct crossed the BBB more efficiently than feRMA1 and feRMA3. This can be attributed to the varying sizes of the GS linker, which potentially affects the efficiency of reverse transcytosis. Further, feRMA expressed in the brain through AAV delivery was rapidly cleared from the serum following the injection of TEV protease. Approximately 86.5%–93.2% of the feRMA signal was quenched within the first 30 min following a single intravenous administration of TEV protease.

The ability to monitor dynamic changes in gene expression in the brain is critical for connecting the behavior and its cellular basis and identification of new drug targets^27,28^. Here, we confirmed that feRMA can be released from various brain regions into the bloodstream with a measurable signal. Our results also demonstrated that feRMA could be rapidly cleared from the blood upon exposure to TEV protease. Additionally, even after signal elimination, the protein levels continued to rise following cleavage, suggesting that the protein continues to be secreted and delivered into the bloodstream owing to the activity of the promoter. We found that repeated feRMA cleavage was possible. We tested the activity of TEV protease in 1 week after feRMA expression and 2 weeks later as well to examine whether immunization against a TEV protein could prevent cleavage. We found that the feRMA baseline signal continued to reduce even after the second administration of TEV protease.

We established that the feRMA approach improves the dynamic range of measurement when paired with weakly expressed promoters, such as the RAM system promoter for measuring the baseline activity of c-Fos in the mouse hippocampus. Relative to the feRMA signal in the vehicle control group, the signal in the TEV protease group increased significantly by 5.51 ± 0.245 fold (means and SEM provided, p < 0.0001, two-tailed unpaired t-test with unequal variance). This result suggests that feRMA accumulates even at moderate levels of neuronal activity, whereas previously, RMA were shown to be useful only after strong chemogenetic activation^6,7,29,30^. Future research could expand on the concept of modifiable serum markers. By proteolytic cleavage, one can reduce the background signal of the long-lasting serum markers. Similarly, by inducing the production of markers in a specific site of the body with a drug and observing the changes in serum, one can infer the underlying cellular signaling of that site. Finally, by binding or chemically modifying the serum markers in vivo, one can also extend their half-life to improve the detection at the cost of temporal resolution, or direct them to more efficiently leave the tissue for easier detection. Overall, in the proof of concept system, feRMA demonstrated sensitivity comparable to that of unmodified RMA but with the added feature of a tunable half-life, making them well-suited for neuroscience applications requiring repeated measurement of gene expression. The paradigm of modifying the markers in vivo to improve their detection is broadly applicable to either natural or synthetic serum markers.

## MATERIALS AND METHODS

### Animals

Wild-type male and female C57BL/6J mice, aged 9–12 weeks, were purchased from Jackson Laboratory (Bar Harbor, ME, USA). Animal experiments were conducted in accordance with the guidelines of the National Institutes of Health and approved by the Institutional Animal Care and Use Committee of Rice University.

### Plasmid construction

To construct pET28a-T7-feRMA-6xHis for protein purification, our previous plasmid pET28a-T7-RMA-6xHis was digested with XcmI and NcoI to obtain its backbone. The feRMA fragment (Gluc-TEVcs-IGg1 FC) purchased from Twist inc was amplified using polymerase chain reaction (PCR ) and inserted into the digested backbone using Gibson assembly. To construct AAV-hSyn-feRMA (Gluc-Fc), the plasmid pAAV-hSyn-Gluc-RMA-IRES-EGFP (Addgene, 189629) was purchased and used as an AAV vector to insert the amplified feRMA fragment into the digested backbone using Gibson assembly.

For AAV-TRE-feRMA-IRES-EGFP, AAV-TRE-RMA-IRES-EGFP was digested with PmeI and BamHI to obtain its backbone and the segment feRMA2 (Gluc-TEVcs-IGg1 FC) was used as an insert for Gibson assembly.

### PC-12 culture for luciferase assay

PC-12 cells (American Type Culture Collection, CRL-1721) were cultured in RPMI 1640 medium (Corning, 10-040-CV) supplemented with 10% horse serum (Life Technologies, 26-050-088) and 5% fetal bovine serum (Corning, 35-011-CV). The cells were incubated in an atmosphere of 5% CO_2_ at 37°C and passaged every 2 days at subculture ratios of 1:2 or 1:3 according to the confluency ratio.

For *in vitro* luciferase assay, PC-12 cells were seeded in 12-well plates at a density of 200,000 cells per well. After 16–20 h, the cells were transfected with 1500 ng of plasmids encoding hSyn-feRMA and 3.0 µL of Lipofectamine 2000 (Life Technologies, 11-668-019), following the manufacturer’s protocol. Culture media samples of 50 µL were collected every 24 h up to 120 h and stored at −20°C. For preparing Gluc substrate, 0.5 mM native coelenterazine stock (Nanolight Technology, 303) containing 66% dimethyl sulfoxide was dissolved in luciferase assay buffer (10 mM Tris, 1 mM EDTA, 1.2 M NaCl, pH 8.0) and stored at −80°C. Before measuring bioluminescence, the coelenterazine stock was diluted to 20 µM in luciferase assay buffer and kept in the dark at 25 °Cor 1 h. The samples were thawed, and 25 µL of each sample was transferred to a black 96-well plate (Corning). An Infinite M Plex microplate reader equipped with i-control software (Tecan) was used to inject 50 µL of assay buffer containing coelenterazine and measure photon emission integrated over 30 s. The measured values were averaged in Excel (Microsoft) to obtain the light units per second.

### Protein purification

Protein purification was performed following the protocol described in our previous study^6^. Briefly, SHuffle T7 Express chemically competent *Escherichia coli* cells (New England Biolabs, C3029J) were transformed with the plasmid pET-T7-feRMA(1,2,3)-His. Approximately 16–20 h post transformation, a single colony was picked and transferred into 5 mL of Luria Bertani broth. This starter culture was grown overnight at 30°C in a shaker incubator shaking at 250 rpm. The culture was then transferred into 1 L of Terrific Broth and grown under the same conditions. The optical density of the culture was monitored at 600 nm until it reached 0.5. Subsequently, feRMA expression was induced using 100 µM isopropyl β-D-thiogalactopyranoside, and the cells were cultured for 20 h at 16°C with shaking at 180 rpm. After harvesting, the cells were resuspended in lysis buffer (300 mM NaCl, 50 mM NaH_2_PO_4_, 10 mM imidazole, 10% glycerol, and pH 8.0) containing ProBlock Gold protease inhibitor (Gold Biotechnology) and lysed in a sonicator (VCX 130, Sonics & Materials). The lysates were centrifuged at 12,000 g for 30 min at 4°C. The supernatants were incubated with Ni-NTA agarose resin (Qiagen, 30210) and loaded onto chromatography columns (Bio-Rad). The resin was washed, and elution was performed using gravity flow. The proteins feRMA1, feRMA2, and feRMA3 were eluted using lysis buffer containing 500 mM imidazole and buffer-exchanged into phosphate-buffered saline (PBS) using Amicon centrifugal filter units with a 10-kDa cutoff (Millipore Sigma). The purity of the proteins in PBS was finally analyzed using sodium dodecyl sulfate–polyacrylamide gel electrophoresis. Protein concentrations were determined using the bicinchoninic acid protein assay (Thermo Fisher Scientific, 23225).

### AAV production

PHP.eB AAV virus was packaged according to the protocol described in our previous study^6^. Briefly, HEK293T cells (American Type Culture Collection, CRL-3216) were transfected with the transfer plasmid, PHP.eB iCap (Addgene, 103005), and pHelper plasmids. After 24 h, the culture medium was replaced with fresh DMEM (Corning, 10-013-CV) containing 5% fetal bovine serum and nonessential amino acids (Life Technologies, 11140050). At 4 days post transfection, the cells were harvested and the culture media were collected. Each sample of culture medium was mixed with a fifth of its volume of polyethylene glycol solution (40% polyethylene glycol 8000, 2.5 M NaCl) to precipitate AAV at 4°C for 2 h. The harvested cells were resuspended in PBS and lysed using the freeze–thaw method. The precipitated AAV was pelleted via centrifugation, resuspended in PBS, combined with the lysed cells, and incubated at 37°C for 45 min.

The AAV viruses were purified using iodixanol gradient ultra-centrifugation. A Quick-Seal tube (Beckman Coulter, 344326) was loaded with iodixanol (Sigma-Aldrich, D1556) gradients of 60%, 40%, 25%, and 15%. After centrifuging the cell lysate, the supernatant was loaded on top of the iodixanol layers. The tube was sealed and centrifuged at 350,000 g for 2.5 h in an ultracentrifuge (Beckman Coulter) using a Type 70 Ti fixed-angle rotor. The AAV viruses were collected by extracting the 40%–60% iodixanol interface and washed in an Amicon centrifugal filter unit with a 100-kDa cutoff (Millipore Sigma). The final AAV virus solution was obtained by filtering through a 0.22-µm polyether sulfone membrane. Viral titers were determined using quantitative PCR.

### Stereotaxic injection

Intracranial injections were achieved using a microliter syringe equipped with a 34-gauge beveled needle (Hamilton) connected to a motorized pump (World Precision Instruments) using a stereotaxic frame (Kopf Instruments). To inject feRMA proteins, feRMA constructs (20 pM) were injected into the CP (AP + 0.25 mm, ML +2.0 mm, DV − 3.2 mm) at a rate of 200 nL min^−1^, and the needle was kept in place for 5 min before taking it out from the injection site. PHP.eB serotype was used in all experiments involving AAV injections. To deliver AAV encoding hSyn-feRMA-IRES-GFP, 2.3 × 10^9^ vp/g in a volume of 200 nL was injected per site at a rate of 600 nL min^−1^ to the following coordinates: CP in the striatum (AP + 0.25 mm, ML + 2.0 mm, DV − 3.2 mm), CA1 in the hippocampus (AP − 1.94 mm, ML + 1.0 mm, DV − 1.3 mm), and substantia nigra in the midbrain (AP − 3.28 mm, ML + 1.5 mm, DV − 4.3 mm).

For chemogenetic neuromodulation, mice were fed 40 mg kg^−1^ Dox chow (Bio-Serv) 48 h before surgery. The following AAV plasmids were used: hSyn-hM3Dq-RAM-d2tTA (2.0 × 10^9^ vp/g) and TRE-Gluc-feRMA-IRES-GFP (2.0 × 10^9^ vp/g). For each surgery, the AAV cocktail was prepared in a volume of 450 nL and injected over 1 min. Dox chow was withdrawn 48 h before blood collection as a baseline to record the neuromodulation.

### Blood collection for luciferase assay

Mice were anesthetized in 1.5%–2% isoflurane in air or O_2_. Subsequently, 1–2 drops of 0.5% ophthalmic proparacaine were applied topically to the cornea of one eye. A heparin-coated microhematocrit capillary tube (Thermo Fisher Scientific, 22-362566) was used for blood collection. The tube was placed into the medial canthus of the eye, and the retro-orbital plexus was punctured to withdraw 50–100 µL of blood. The blood samples were centrifuged twice at 1,500 g for 10 min to separate the plasma. For the luciferase assay, 5 µL of plasma was mixed with 45 µL of PBS + 0.001% Tween 20 in a black 96-well plate. The bioluminescence of Gluc-feRMA was measured using a microplate reader after injecting 50 µL of 20 µM coelenterazine dissolved in luciferase assay buffer.

### Histological imaging and analysis

Mice brains were extracted and fixed overnight in 10% neutral-buffered formalin (Sigma-Aldrich, HT501128). Coronal sections were cut at a thickness of 50 µm using a vibratome (Leica), stained in blocking buffer (0.2% Triton X-100 and 10% normal donkey serum in PBS) for 2 h at room temperature, probed overnight at 4°C with primary antibodies, washed thrice in PBS for 15 min, and incubated for 4 h at room temperature with secondary antibody. After a final few washes in PBS, the sections were mounted on glass slides using the mounting medium (Vector Laboratories). The antibodies used were as follows: rabbit anti-Gluc (1:1500, Nanolight Technology) and Alexa 594 secondary antibody (1:500, Life Technologies).

Images were captured using a BZ-X800 fluorescence microscope (Keyence). Manual cell counting was performed using ZEN Blue software (Zeiss) by a blind observer who was not informed about the experimental conditions. Cells in all other images were counted individually.

### Intravenous injection of proteins and TEV protease

Mice were anesthetized. A catheter equipped with a 30-gauge needle was used for mouse tail vein injections. After penetrating along the mouse vein, the catheter was secured in place using tissue glue. FeRMA was injected through the catheter at a dose of 20 pM and blood was collected at timepoints of 2, 6, 12, and 24 h. At 2 h post blood collection, TEV protease was injected at different volumetric doses of 5, 16, and 160 µl. The mice were euthanized after 24 h and their brains were extracted for histological analysis. For subsequent experiments, TEV protease was administered at a volumetric dose of 160 μL.

### Statistical analysis

Means between more than two datasets were compared using one-way ANOVA with Tukey’s honestly significant difference post hoc test. Two-tailed unpaired t-test with unequal variance was used to compare two datasets. Two-way ANOVA with Sidak’s multiple comparison tests were used to compare datasets with two or more variables. P values were determined using Prism (GraphPad Software), and the following notation was adopted throughout the manuscript: not significant, P ≥ 0.05; *P < 0.05; **P < 0.01; ***P < 0.001; and ****P < 0.0001. P values and statistical test results are available in the source data. Figures were made using Adobe Illustrator. A full mixed-effects model was constructed using JMP Pro software (SAS Institute Inc.) to assess the impact of TEV protease on the fold-change in RMA levels in blood serum across three brain regions: the hippocampus, dopaminergic midbrain, and striatum. The model included as fixed effects whether RMA levels were assessed before or after TEV administration (TEV or vehicle) and the number of weeks (1 or 3) since injection of the designed AAV. Each mouse served as a biological replicate, and mice were modeled as random effects to account for between-subject variability in both the slope and intercept of the modeled effects. Separate models were fitted for each brain region. This approach allowed for robust estimation of fixed effects while capturing individual differences in baseline levels and trajectories across animals. The same model was also used to measure changes in the overall feRMA levels over time and the effects of TEV protease during the *in vivo* measurement of baseline c-Fos activity.

Male C57BL/6J mice between 10 and 14 weeks old were obtained from Jackson Laboratory (Bar Harbor, ME, USA). Animals were housed in a 12-hour light/dark cycle and were provided with water and food ad libitum. All experiments were conducted under a protocol approved by the Institutional Animal Care and Use Committee (IACUC) of Rice University.

## Data availability

The authors declare that all data supporting the results in this study are available within the paper, its Supplementary Information and its Source Data file. Microscopy images are available from the corresponding author upon reasonable request owing to their large size and numbers.

## ACKNOWLEDGEMENTS

The work described in this study was supported by a grant from the NIH (DP2EB035905) to J.O.S, and by the NSF GRFP fellowship to E.K.R.

## AUTHOR CONTRIBUTIONS

J.O.S. and S.N. conceived and planned the research. S.N. and J.O.S. designed the experiments and wrote the paper, with input from all other authors. S.N. performed and participated in all experiments described in the study. S.L. constructed and designed the plasmids. S.L. and V.A.S. performed in vitro experiments. H.L assisted with retro-orbital blood collection and TEV protease administration. H.L. and E.K.R. performed histological processing and counting. V.P.C., J.O.S., and S.N. performed data analysis. J.O.S. supervised the study.

