## supplementary data for "Erasable Serum Markers"

### Erasable Synthetic Serum Markers

**SUPPLEMENTARY FIGURES AND TABELES:**

#### Table1 FERMA sequences:

**Supplemantry table 1. feRMA 1-3 sequences.** Highlighted sequence (yellow) corresponds to the TEV cleavage site with the added linker sequences.

##### Modification of pAAV-hSyn-Gluc-RMA-IRES-EGFP

(Plasmid #189629)

| | |
|---|---|
| feRMA construct1 | ggagatataccatgggcagcagccatcatcatcatcatcacagcagcggcaagcccaccga-<br>gaacaacgaagacttcaacatcgtggccgtggccagcaacttcgcgaccacggatctcgatgctgac<br>cgcgggaagttgcccggcgagaagctgccgctggaggtgctcaaagagttggaa-<br>gccaatgcccggaaagctggctgcaccaggggctgtctgatctgcctgtcccacatcaagtgcacgc<br>ccaagatgaagaagttcatcccaggacgctgccacacctacgaaggcgacaaagagtccg-<br>cacagggcggcataggcgaggcgatcgacgacattcctgagattcctgggttcaaggacttggagcc<br>catagagcagttcatcgcacaggtcgatctgtgtgtggactg-<br>cacaactggctgcctcaaagggcttgccaacgtgcagtgttctgacctgctcaagaagtggctgccgc<br>aacgctgcgcgacctttgccagcaagatccagggccaggtggacaa-<br>gatcaagggggccggtgatgacGGCTCCGAGAACCTGTACTTCCAGTCCgt<br>cccagaagtatcatctgtcttcatcttcccccccaaagcccaaggatgtgctcac-<br>cattactctgactcctaaggtcacgtgtgttgtggtagacatcagcaaggatgatcccgaggtccagttc |

| | |
|---|---|
| | agctggtttgtagatgatgtggaggtgcacacagctcagacgcaaccccgggaggagcag-<br>ttcaacagcactttccgctcagtcagtgaacttcccatcatgcaccaggactggctcaatggcaaggag<br>ttcaaatgcagggtcaacagtgcagctttccctgcccccatcgagaaaaccatctccaaaac-<br>caaaggcagaccgaaggctccacaggtgtacacc |
| feRMA construct 2 | ggagatataccatgggcagcagccatcatcatcatcatcacagcagcggcaagcccaccga-<br>gaacaacgaagacttcaacatcgtggccgtggccagcaacttcgcgaccacggatctcgatgctgac<br>cgcgggaagttgcccggcgagaagctgccgctggaggtgctcaaagagttggaa-<br>gccaatgcccggaaagctggctgcaccaggggctgtctgatctgcctgtcccacatcaagtgcacgc<br>ccaagatgaagaagttcatcccaggacgctgccacacctacgaaggcgacaaagagtccg-<br>cacagggcggcataggcgaggcgatcgacgacattcctgagattcctgggttcaaggacttggagcc<br>catagagcagttcatcgcacaggtcgatctgtgtgtggactg-<br>cacaactggctgcctcaaagggcttgccaacgtgcagtgttctgacctgctcaagaagtggctgccgc<br>aacgctgcgcgacctttgccagcaagatccagggccaggtggacaa-<br>gatcaaggggggccggtgatgacGGCTCCGAGAACCTGTACTTCCAGTCCG<br>GTTCAgtcccagaagtatcatctgtcttcatcttccccccaaagcccaaggatgtgctcac-<br>cattactctgactcctaaggtcacgtgtgttgtggtagacatcagcaaggatgatcccgaggtccagttc<br>agctggtttgtagatgatgtggaggtgcacacagctcagacgcaaccccgggaggagcag-<br>ttcaacagcactttccgctcagtcagtgaacttcccatcatgcaccaggactggctcaatggcaaggag<br>ttcaaatgcagggtcaacagtgcagctttccctgcccccatcgagaaaaccatctccaaaac-<br>caaaggcagaccgaaggctccacaggtgtacaccattcc |

| | |
|---|---|
| feRMA construct 3 | Atgggcagcagccatcatcatcatcatcacagcagcggcaagcccaccgagaacaacgaa-<br>gacttcaacatcgtggccgtggccagcaacttcgcgaccacggatctcgatgctgaccgcgggaagt<br>tgcccggcgagaagctgccgctggaggtgctcaaagagttggaagccaatgcccggaaa-<br>gctggctgcaccaggggctgtctgatctgcctgtcccacatcaagtgcacgcccaagatgaagaagtt<br>catcccaggacgctgccacacctacgaaggcgacaaagagtccgcacagggcgg-<br>cataggcgaggcgatcgacgacattcctgagattcctgggttcaaggacttggagcccatagagcagt<br>tcatcgcacaggtcgatctgtgtgtggactgcacaactggctgcctcaaagggcttgccaac-<br>gtgcagtgttctgacctgctcaagaagtggctgccgcaacgctgcgcgacctttgccagcaagatcca<br>gggccaggtggacaagatcaagggggccggtgatgacGGCTCCGAGAACCT-<br>TACTTCCATCCGGTTCAGGTGGTTCTgtcccagaagtatcatctgtcttcatcttc<br>cccccaaagcccaaggatgtgctcaccattactctgactcctaaggtcacgtgtgttgtgg-<br>tagacatcagcaaggatgatcccgaggtccagttcagctggtttgtagatgatgtggaggtgcacaca<br>gctcagacgcaaccccgggaggagcagttcaacagcactttccgctcagtcag-<br>tgaacttcccatcatgcaccaggactggctcaatggcaaggagttcaaatgcagggtcaacagtgcag<br>ctttccctgcccccatcgagaaaaccatctccaaaaccaaaggcagac-<br>cgaaggctccacaggtgtacaccattccacctcccaaggagcagatggccaaggataaagtcagtct<br>gacctgcatgataacagacttcttccctgaagacattactgtggagtggcagtggaatggg-<br>cagccagcggagaactacaagaacactcagcccatcatggacacagatggctcttacttcgtctacag<br>caagctcaatgtgcagaagagcaactgggaggcaggaaatactttcac-<br>ctgctctgtgttacatgagggcctgcacaaccaccatactgagaagagcctctcccactctcctggtaa<br>atgag |

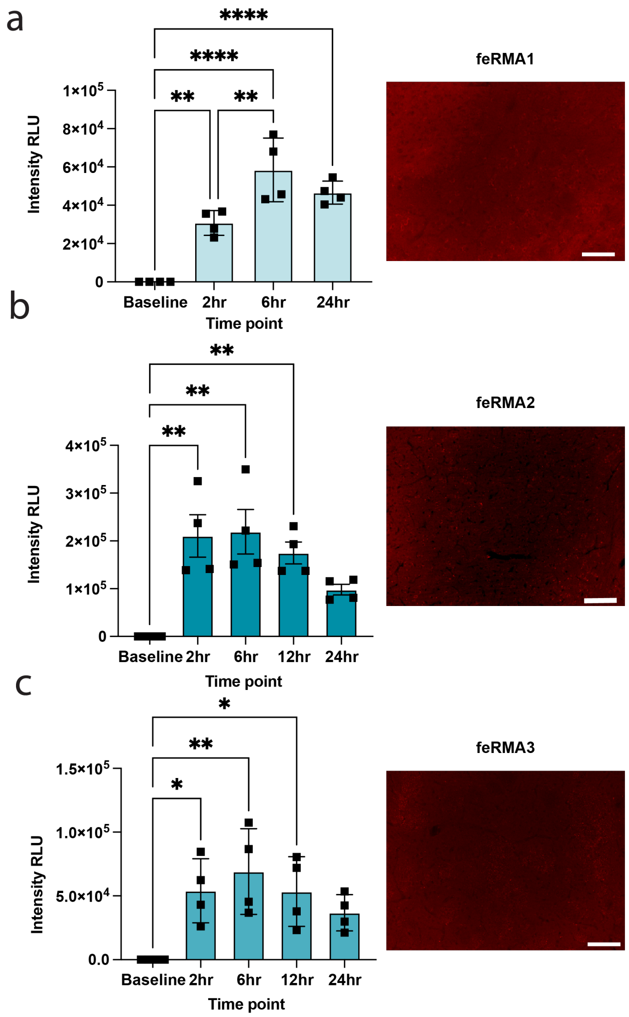


**Supplementary Figure S1. feRMA protein constructs evaluation in vivo**. Plasma bioluminescence signal and 20x representative images of CP to evaluate feRMA expression in vivo at different time point for a, feRMA1 b, feRMA2 c, feRMA3. N = 4 independent mice were analyzed. In comparison with the signal at 0 h, using one-way ANOVA, with Tukey's test: in **a**, $P$ = 0.0028 (0 h versus 2 h) and $P < 0.0001$ (0 h versus 6 h) ($F_{3,12} = 28.08$, $P < 0.0001$). in **b**, $P = 0.0018$ (0 h versus 2 h) and $P = 0.0013$ (0 h versus 6 h) ($F_{4,15} = 8.818$, $P = 0.0007$). in **c**, $P = 0.0347$ (0 h versus 2 h) and $P = 0.0059$ (0 h versus 6 h) ($F_{4,15} = 5.164$, $P = 0.0081$). Means with standard error of the mean (SEM) presented in bar graphs. Scale bar 100 μm

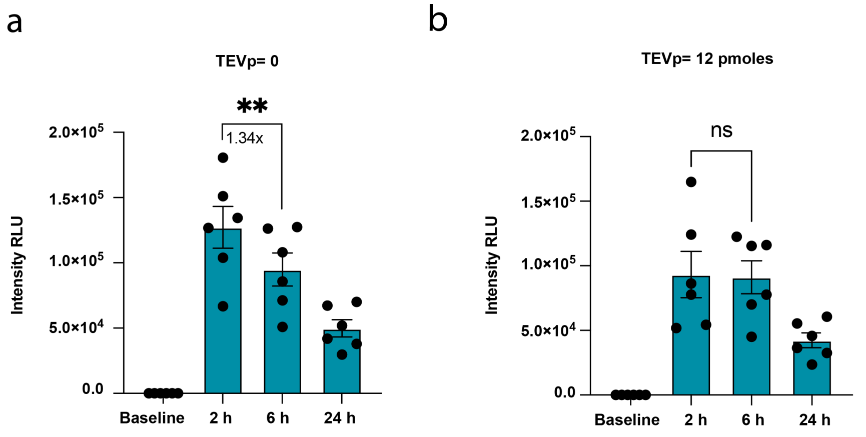


**Supplementary Figure S2. Optimization of TEV protease dose to enable feRMA cleavage *in vivo*.** A total of 20 pmols of feRMA were injected intracranially into the CP of the brain, and TEV protease was injected at different doses to evaluate the cleavage efficacy. Plasma bioluminescence signal was measured at each relevant TEV protease dose. The signal at 2 h was compared with that at 6 h using one-way ANOVA with Sidak's test. **a**, n = 6 (TEV = 0 µg) independent mice were analyzed. P = 0.0022 (($F_{1.161,5.804} = 57.99$, $P = 0.0003$), 2 h vs. 6 h at 0 dose). **b**, n = 6 (TEV = 12 pmols) independent mice were analyzed, P = 0.8844 (($F_{1.664,8.319} = 23.76$, $P = 0.0005$), 2 h vs. 6 h at 12 pmols dose). ns (not significant, p > 0.05), **P < 0.01. Means with SEM presented in bar graphs.

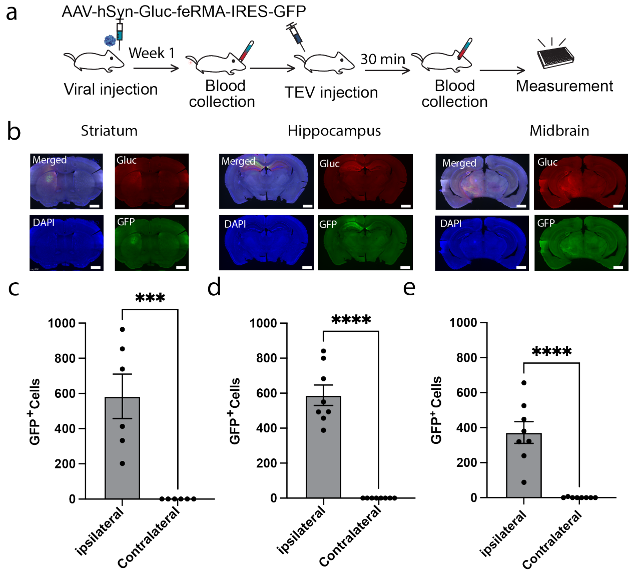


**Supplementary Figure S3. a,** Experimental scheme for detecting gene expression following AAV injection in the brain using feRMA. **b**, Representative 4× images showing expression of feRMA in the target brain regions (CP, CA1, and substantia nigra), respectively. Scale, 1000 µm. **c**-**e**, Quantification of GFP expression at the ipsilateral in images represented in b. n = 6 (CP) and n = 8 (CA1 and substantia nigra) independent images were analyzed, P = 0.001 (CP GFP+; t = 4.620, df = 10), P < 0.0001 (CA1, GFP+; t = 9.986, df = 14), and P < 0.0001 (SN GFP+; t = 6.018, df = 14), using two-tailed unpaired t-test. ***P < 0.001, ****P < 0.0001. Means with SEM presented in bar graphs.
